# Three invariant Hi-C interaction patterns: applications to genome assembly

**DOI:** 10.1101/306076

**Authors:** Sivan Oddes, Aviv Zelig, Noam Kaplan

**Affiliations:** Department of Physiology, Biophysics & Systems Biology, Rappaport Faculty of Medicine, Technion - Israel Institute of Technology, Haifa, Israel

**Keywords:** Computational biology, Genomics, Genome assembly, Genome scaffolding, 3D genome, Hi-C

## Abstract

Assembly of reference-quality genomes from next-generation sequencing data is a key challenge in genomics. Recently, we and others have shown that Hi-C data can be used to address several outstanding challenges in the field of genome assembly. This principle has since been developed in academia and industry, and has been used in the assembly of several major genomes. In this paper, we explore the central principles underlying Hi-C-based assembly approaches, by quantitatively defining and characterizing three invariant Hi-C interaction patterns on which these approaches can build: Intrachromosomal interaction enrichment, distance-dependent interaction decay and local interaction smoothness. Specifically, we evaluate to what degree each invariant pattern holds on a single locus level in different species, cell types and Hi-C map resolutions. We find that these patterns are generally consistent across species and cell types but are affected by sequencing depth, and that matrix balancing improves consistency of loci with all three invariant patterns. Finally, we overview current Hi-C-based assembly approaches in light of these invariant patterns and demonstrate how local interaction smoothness can be used to easily detect scaffolding errors in extremely sparse Hi-C maps. We suggest that simultaneously considering all three invariant patterns may lead to better Hi-C-based genome assembly methods.

## 1. Introduction

Since the publication of the draft human genome sequence, advances in DNA sequencing technology have transformed modern biological research [1]. High throughput next-generation sequencing (NGS) technology, based on short reads, has allowed to obtain massive amounts of genomic sequences. In spite of this, standard short read NGS technology alone is not sufficient to produce reference-quality genomes[2,3]. Ironically, the substantial ease of NGS compared to traditional low-throughput methods (e.g. involving cloning), has led to a stark decrease in the quality of published genomes, when compared to traditionally-sequenced genomes such as that of human and mouse.

To understand the problems associated with standard short-read NGS assembly, let us briefly overview how NGS is typically used for *de novo* genome assembly. First, genomic DNA is fragmented and sequenced. Resulting reads that contain unique overlapping sequences are then stitched together to create longer contiguous sequences called *contigs*. Due to the repetitive nature of genomes and their size relative to the read size, many overlaps will be nonunique, resulting in a huge number of contigs (on the order of 10^5^-10^6^ contigs for a human-sized genome). Next, contigs are grouped and positioned relative to each other in a process called *scaffolding*. Typically, long-insert libraries are used for scaffolding. This consists of genome fragmentation, size-selection to a predetermined size range and paired-end sequencing. Molecules that uniquely map to two different contigs can then be used to estimate the distance between the contigs and position them relative to one another. Each resulting set of associated contigs (including gaps in between) is called a *scaffold*. Ideally, one would like to obtain a single scaffold for each chromosome. However, even with high coverage and multiple-sized long-insert libraries, a human-sized genome may end up having ~10^4^-10^5^ scaffolds. While these highly fragmented genomes may be useful for some applications, they have limited utility for the study of long-range phenomena, including large-scale genome evolution, comparative genomics, gene regulation, haplotyping and 3D genome organization. Furthermore, short read NGS cannot accurately reconstruct complex polyploid and rearranged genomes - including cancer genomes. Thus, these limitations seriously hinder our genomic view of many critical biological systems.

Importantly, the dramatic decrease of cost per read produced by NGS will not help alleviate problems associated with non-uniqueness due to read length. The recent development of long-read sequencing technologies, has helped mitigate some of the problems associated with short reads and can reduce the number of scaffolds by one or two orders of magnitude, but is generally expensive and limited in achieving chromosome-scale scaffolds [4,5]. Alternatively, non-sequencing based technologies exist, ranging from classical cloning to newer techniques such as optical mapping [6,7]. Thus, the development of novel simple techniques that can take advantage of short read sequencing remains an important challenge.

Hi-C is a molecular biology technique which measures spatial physical proximity between pairs of DNA loci genome-wide [8,9]. Hi-C is based on proximity ligation, such that DNA sequences that are in physical proximity are ligated to each other, and are then measured with NGS technology. By sequencing hundreds of millions of these chimeric molecules, data can be aggregated to construct an interaction matrix which provides an interaction frequency for every pair of genomic loci. Interaction patterns observed in the interaction map are then interpreted and used to extract biological knowledge. Hi-C and similar techniques [10,11], all of which are derivatives of Chromosome Conformation Capture (3C) [12], has been used extensively to study genome 3D organization and led to key biological insights in several different species, including bacteria [13], yeast [14,15], plants [16], worms [17], insects [18] and mammals [8,19–22]. Although in this paper we focus on Hi-C, most of this work is relevant for other genome-wide 3C derivatives.

Recently, we and others have proposed that Hi-C measurements can be used to solve outstanding challenges related to genome assembly [23–33]. This is based on the notion that all Hi-C interaction maps share common features which relate 3D interaction frequencies to the 1D ordering of the genome. For example, loci which are nearby in the genomic sequence tend, on average, to interact more frequently than loci that are located far away on the chromosome, which in turn interact more frequently than loci that are located on different chromosomes [23,34]. Thus, given a set of contigs and Hi-C measurements on a new genome, we can build on these shared principles, consolidating interaction frequencies between contigs to estimate the relative positions of contigs and scaffold them. Advantages of this approach include robustness to very large gaps, relatively low sequencing coverage requirement and applicability to any species. Additionally, a major feature of this approach is that it is based on short-read technology, and can thus directly benefit from the rapid increase in short-read sequencing power. Since the initial proposal of this notion [23,24], Hi-C has been used to scaffold several major genomes including the frog [35], quinoa [36], goat [37], mosquito [29], barley [38], house spider [39], alligator [40], durian [41], lettuce [42] and cassava [42] genomes. Still, Hi-C is typically used for scaffolding in conjunction with intermediate techniques, such as long-read sequencing.

Hi-C is also useful for addressing other challenges related to genome assembly. (1) <u>Haplotype phasing:</u> Large-scale haplotype phasing is difficult with short reads, since Single Nucleotide Polymorphisms (SNPs) may be sparse and only reads that map to two SNPs are useful. Due to the physical separation of homologous chromosomes in the nucleus, the probability of observing intrachromosomal interactions is much higher than observing interchromosomal interactions. Thus, Hi-C can be used to reconstruct chromosome-scale scaffolds by providing SNP linkage information over large distances that are not spanned by normal reads [31,43]. (2) <u>Metagenome deconvolution:</u> In metagenome and microbiome samples, sequencing typically results in a set of contigs and the challenge is to group contigs that come from the same species. Hi-C provides long-distance linkage information and can thus be used to connect contigs, while the problem of observing genomic interactions between cells is extremely low [26–28]. Hi-C can also be used to associate plasmids with their respective genomes. (3) <u>Cancer genomes:</u> Cancer genomes are often highly rearranged and thus pose a major challenge to assemble *de novo*. Large structural variations are especially difficult to measure with short reads, since the rearrangement will only be reflected in a small fraction of the reads (i.e. those that map to the edges of the rearranged region). In Hi-C, structural variations can be detected since they appear to deviate from standard Hi-C patterns [32,33].

## 2. Invariant Hi-C patterns

Since the 3D organization of a genome reflects its functional state, it is not surprising that 3D genome organization differs between species. In fact, 3D genome organization varies between cell-types [8,20], along different stages of the cell-cycle [44–46], and even within homogenous populations of synchronized cells [46]. Despite this, certain aspects of 3D genome organization, as measured by Hi-C, are universal [23,34]. We refer to these as *invariant patterns*, since they have been observed across species, cell types and conditions. These patterns have even observed in single-cell interaction maps [46,47]. Importantly, other biological patterns, which are specific to the biological system at hand, will lead to local deviations from these general rules, but by large the invariant patterns are dominant in Hi-C data. In fact, these patterns are so robust and ubiquitous that they are used to evaluate the quality of Hi-C experiments and check for experimental artifacts [34]. Importantly, these patterns have previously been studied quantitatively mainly at the level of genome averages, but less at the level of individual loci. In this paper, we set out to explore the central principles underlying Hi-C-based assembly, by quantitatively defining and characterizing three invariant Hi-C interaction patterns on which these approaches can build: Intrachromosomal interaction enrichment, distance-dependent interaction decay and local interaction smoothness. Specifically, we evaluate to what degree each invariant pattern holds on a single locus level in different species, cell types and Hi-C map resolutions/sequencing depths. Evaluation at the single-locus level is important since loci which deviate from these patterns may lead to assembly errors. We note that a general limitation of these analyses is that Hi-C does not directly measure interaction probabilities, meaning that due to the random sampling and sequencing-depth limitation inherent to the experiment, the estimation of small interaction probabilities will be unreliable.

In our analyses we use Hi-C maps from human (Hap1 [48], IMR90 [19], HESC [19]), mouse (MESC [19], cortex [19]), worm (*C. elegans* [17]) and bacteria (*C. crescentus* [13]). All maps were processed using the Dekker lab Hi-C pipeline and balanced unless specified otherwise [49,50]. All code required to reproduce the results in this paper is available at https://github.com/KaplanLab/Invariants.

### 2.1 Invariant pattern I: Intrachromosomal interaction enrichment

The first invariant pattern is intrachromosomal interaction enrichment. In Hi-C interaction maps, this is observed as a tendency of loci to interact more frequently with loci within the same chromosome (cis-interactions) than with loci on different chromosomes (trans-interactions). Two major components underlie this pattern. The first component is a phenomenon known as *chromosome territories*, in which chromosomes occupy distinct volumes throughout cell cycle, leading to physical separation between chromosomes [51]. The second component is the random positioning of chromosomes in the nucleus. Although some chromosomes show a tendency to be located near the center of the nucleus while others tend to be located more peripherally (this is known as radial positioning), the relative positions of chromosomes with respect to each other is largely random [52]. This may be due to the inability of relative chromosome positioning to be inherited through cell-cycle, which could lead to such variation in a cell population [53]. Thus, on a population average, as measured in a standard Hi-C experiment, the probability of any specific pair of chromosomes to interact over the entire population is low. Notably, this phenomenon may not hold in a singlecell Hi-C interaction map, but chromosome territories will be present in single-cell interaction maps[46,47]. The combination of these two components yields a strong bias towards intrachromosomal interactions in Hi-C interaction maps.

This invariant is explicitly used to evaluate the quality of Hi-C libraries. Typically the ratio between intrachromosomal (cis) and interchromosomal (trans) is used as a quality metric for Hi-C, but, depending on how it is calculated, may be genome-specific as it can depend on the number and sizes of chromosomes. The underlying logic is that general random noise (such as that caused by background ligation) will affect the interaction matrix uniformly, and thus cause cis and trans to be similar [49].

Formally, we can characterize this invariant pattern as:

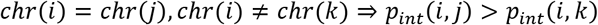

Where *chr*(*i*) is the chromosome of locus *i*, and *P_int_*(*i,j*) is the interaction probability of loci *i,j*. In other words, if *j* is on the same chromosome as *i* but *k* is not, *i* will interact more frequently with *j* than with *k*.

To quantify the extent to which invariant I holds, we used the following scheme. For each genomic locus (i.e. bin) *i* in a Hi-C matrix, we considered all pairs of loci *j,k* such that *j* is on the same chromosome and *k* is on a different chromosome. We then calculated in what fraction of pairs *j* interacts more frequently with *i* than *k* does and refer to this as the *consistency* of locus *i* with invariant I (marked *C_I_*(*i*)). Similarly, we calculated in what fraction of pairs *j* interacts less frequently with *i* than *k* does and refer to this as the *inconsistency* of locus *i* with invariant I (marked 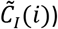.

We then tested the distribution of consistency and inconsistency with invariant I genome-wide in various Hi-C map resolutions, cell types, species and with/without map balancing (Figure 1). We find high median *C_I_* values of ^~^0.99 in 1Mb resolution Hi-C maps across multiple cell types in human (Hap1, HESC, IMR90) and mouse (cortex, MESC). We noted that loci with low consistency are often located near telomeres or centromeres, possibly due to telomere/centromere interchromosomal interactions, low interaction probability at large genomics distances, or errors related to sequence complexity in these regions. For 100Kb maps, we observe that their low read coverage (on average 100 times less reads per matrix entry relative to the 1Mb maps) reduces the median *C_I_* 0.76-0.78. We also observe that Hi-C matrix balancing [49,50] improves the median *C_I_* (0.99/0.96 for balanced/unbalanced Hap1 1Mb; 0.76/0.69 for balanced/unbalanced Hap1 100Kb). Finally, we observe slightly lower median *C_I_* values of 0.80 (50Kb bins) and 0.50 (10 Kb bins) for *C. elegans* Hi-C, which we attribute to both lower sequencing depth and relatively low quality (high noise) of this Hi-C map. In summary, we find that invariant I consistency is high across species and cell types, but can be reduced by low read coverage per bin or by avoiding map balancing.

**Figure 1.**
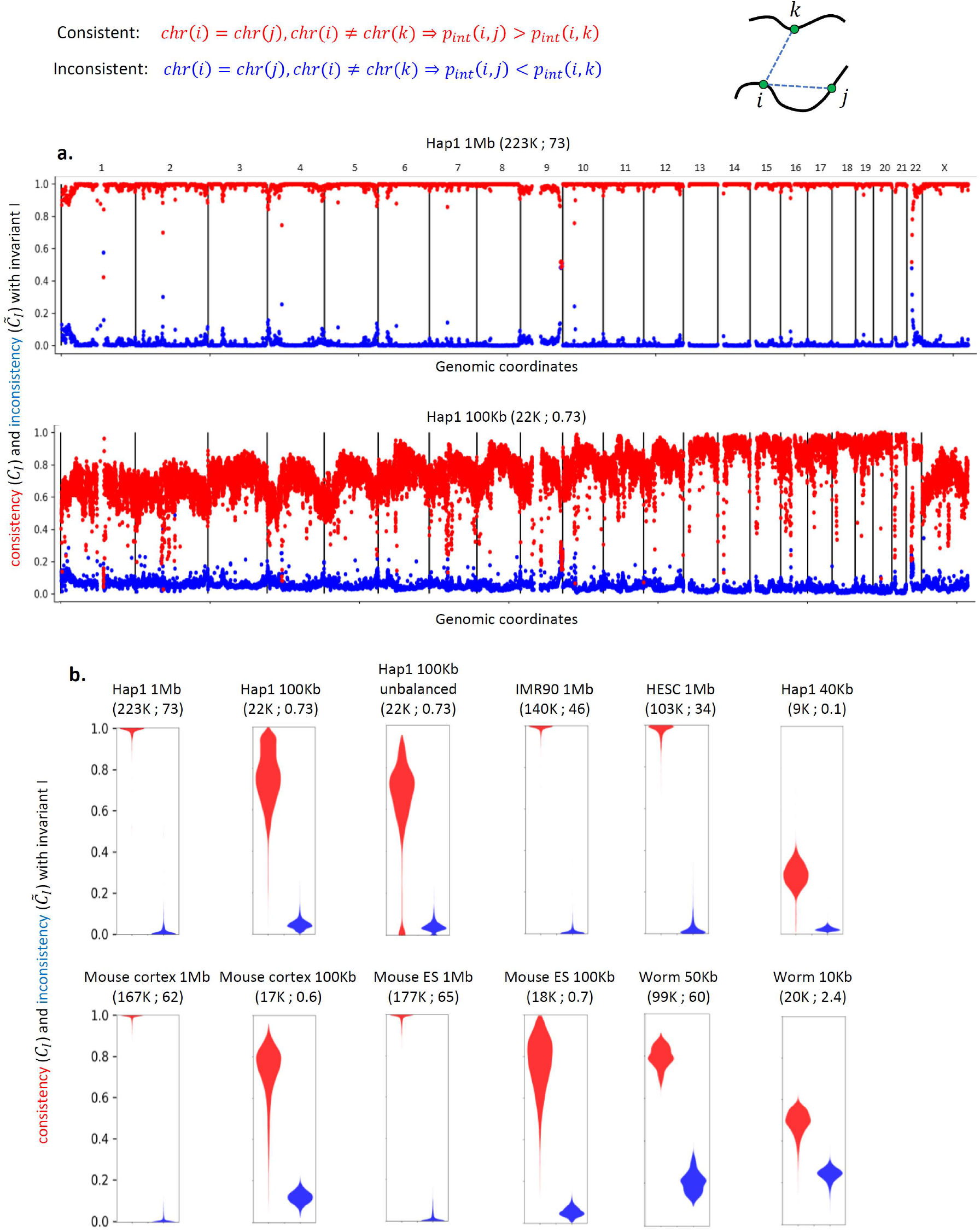
Genome-wide analysis of invariant pattern I (intrachromosomal interaction enrichment). The resolution (bin-size) of each Hi-C map is specified along with the number of valid read-pairs per bin and per matrix entry in parentheses. (**a**) Genome-wide consistency (*C_I_* in red, see text for details) and inconsistency (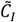 in blue) with invariant I for Hap1 1Mb and 100Kb Hi-C interaction maps. (**b**) Violin plots showing the genome-wide distribution of *C_I_* and 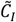 in various Hi-C interaction maps.

### 2.2 Invariant pattern II: Distance-dependent interaction decay

The second invariant pattern is distance-dependent interaction decay. In Hi-C interaction maps, this is observed as a general tendency of interaction frequency to decrease with genomic distance, such that a locus interacts more frequently with loci which are nearby in the genomic sequence than with far away loci. This type of distance-dependent decay is an inherent feature of many polymer physics models, which are inherently stochastic, simply because loci which are nearby in the genomic sequence will interact frequently by random. Thus, we expect this invariant to hold simply due to the stochastic nature of the DNA polymer. In fact, the exact details of the distance dependence can be used to suggest properties of the underlying polymer and it may thus also hold biological relevance, such as in the case of mitotic chromosome organization [45].

Due to the dominance of this invariant, it is often normalized out of the data in order to detect subtler biological patterns such as genomic compartments [8,49,50].

Formally, we can characterize this invariant pattern as:

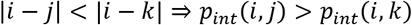

Where *P_int_* (*i,j*) is the interaction probability of loci *i,j*. In other words, if *j* is closer (in genomic distance) to *i* than *k* is, *i* will interact more frequently with *j*. Note that while their underlying mechanisms may be different, invariant II implies invariant I if we define that being on a different chromosome is equivalent to being infinitely far in genomic sequence.

To quantify the extent to which invariant II holds, we used the following scheme. For each genomic locus (i.e. bin) *i* in a Hi-C matrix, we considered all pairs of loci *j, k* such that *j* is closer to *i* in genomic distance than *k* is. We then calculated in what fraction of pairs *j* interacts more frequently with *i* than *k* does and refer to this as the *consistency* of locus *i* with invariant II (marked *C_II_*(*i*)). Similarly, we calculated in what fraction of pairs *j* interacts less frequently with *i* than *k* does and refer to this as the *inconsistency* of locus *i* with invariant II (marked 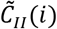).

Next, we tested the distribution of consistency and inconsistency with invariant II genome-wide in various Hi-C map resolutions, cell types, species and with/without map balancing (Figure 2). We find median *C_II_* values in the range of 0.79-0.88 in 1Mb resolution Hi-C maps across multiple cell types in human (Hap1, HESC, IMR90) and mouse (cortex, MESC). In contrast to invariant I, here we find variation which is not necessarily associated with read coverage (e.g. median *C_II_* 0.88 vs 0.79 for mouse ESC vs cortex with roughly similar amount of read coverage). Once again, we noted that loci with low consistency are often located near telomeres or centromeres, likely due to the problematic nature of these regions. For 100Kb maps, we observe lower median *C_II_* values in the range of 0.63-0.74. We also observe that Hi-C matrix balancing improves the median *C_II_* (0.85/0.80 for balanced/unbalanced Hap1 1Mb; 0.73/0.65 for balanced/unbalanced Hap1 100Kb). We observe median *C_II_* values of 0.82 (50Kb bins) and 0.56 (10 Kb bins) for *C. elegans* Hi-C, values which are comparable to the mammalian Hi-C maps when controlling for read coverage. Finally, we tested a bacterial (*C. crescentus;* 10Kb) Hi-C map and found a bimodal *C_II_* distribution (*C_II_* peak centers at 0.53 and 0.82), reflecting the circular nature of the chromosome and highlighting the potential difficulty in applying this principle to circular chromosomes. In summary, we find that invariant II consistency is high, but can vary, across cell types and species, and can be reduced by low read coverage per bin or by avoiding map balancing.

**Figure 2.**
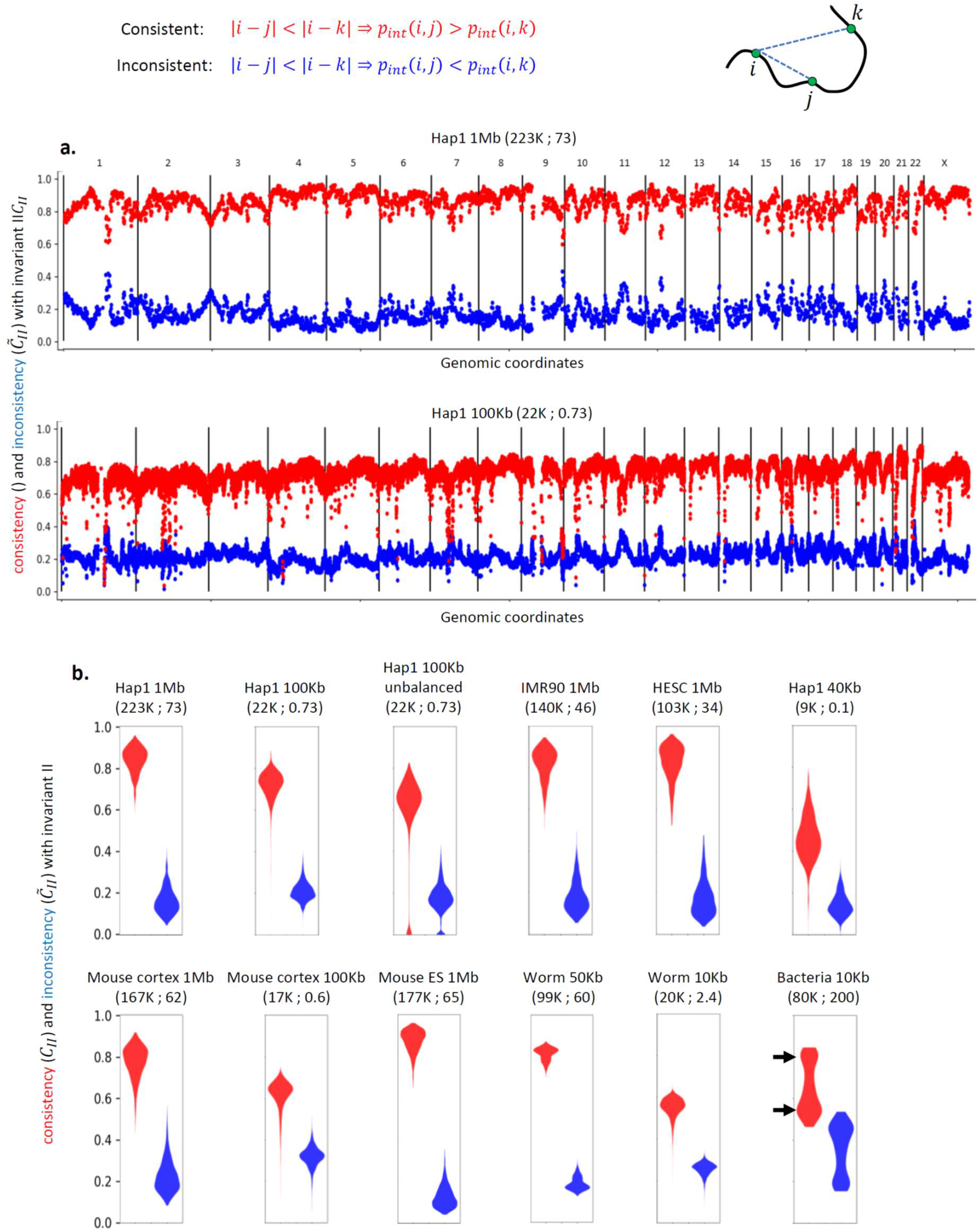
Genome-wide analysis of invariant pattern II (distance dependent interaction decay). (**a**) Genome-wide consistency (*C_II_* in red, see text for details) and inconsistency (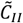 in blue) with invariant II for Hap1 1Mb and 100Kb Hi-C interaction maps. (**b**) Violin plots showing the genome-wide distribution of *C_II_* and 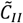 in various Hi-C interaction maps.

### 2.3 Invariant pattern III: Local interaction smoothness

The third invariant is local interaction smoothness. In Hi-C interaction maps, this is observed as a tendency of nearby loci to share similar interactions (e.g. adjacent rows will tend to be similar). The physical basis of this invariant is intuitive: given the set of interactions of a genomic locus A, if we consider a locus B that is sufficiently close to A it will also be close to the neighbors of A simply due to physical constraints such as the triangle inequality and chromatin persistence length. This invariant is informally used to assess experimental artifacts in Hi-C experiments, and one of the criteria for successful removal of biases (e.g. by matrix balancing) is the visual smoothness of the resulting matrix [49,50].

Formally, we can characterize this invariant pattern as:

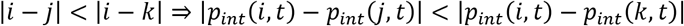

Where *P_int_* (*i,j*) is the interaction probability of loci *i,j*. In other words, if *j* is closer to *i* than *k* is, we expect the interaction probability of (*j, t*) to be closer to that of (*i, t*) than the interaction probability of (*k, t*) is to that of (*i, t*). We expect this to be mostly a local effect, so that it would hold only for *j* which is relatively close to *i*.

To quantify the extent to which invariant III holds, we used the following scheme. For each genomic locus (i.e. bin) *i* in a Hi-C matrix, we considered positions *j = i + 1,k = i + 10* (units are bins) as well as every same-chromosome locus *t*. We calculated for which fraction of *t* the interaction of (*j,t*) is closer (absolute value of the difference) to that of (*i, t*) than the interaction of (*k, t*) is to that of (*i, t*). We refer to this as the *consistency* of locus *i* with invariant III (marked *C_III_*(*i*)). Similarly, we calculated for which fraction of *t* the interaction of (*k, t*) is closer (absolute value of the difference) to that of (*i, t*) than the interaction of (*j, t*) is to that of (*i, t*), and refer to this as the *inconsistency* of locus *i* with invariant II (marked 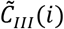). We excluded the main diagonal from this analysis.

We tested the distribution of consistency and inconsistency with invariant III genome-wide in various Hi-C map resolutions, cell types, species and with/without map balancing (Figure 3). We find median *C_III_* values in the range of 0.75-0.79 in 1Mb resolution Hi-C maps across multiple cell types in human (Hap1, HESC, IMR90) and mouse (cortex, MESC). For 100Kb maps, we observe low median *C_III_* values of 0.49. For Hap1 40Kb we observed a very low median *C_III_* value of 0.27 which is only slightly higher than the median 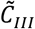 value of 0.25. We find that balancing improves the median *C_III_* (0.79/0.73 for balanced/unbalanced Hap1 1Mb; 0.49/0.38 for balanced/unbalanced Hap1 100Kb. For the bacterial Hi-C map, we observe a high level of consistency (median *C_III_* 0.95 for 40Kb bins and 0.73 for 10Kb bins), suggesting this invariant may alleviate some of the issues associated with circular chromosomes and invariant II. In summary, we find that invariant III consistency is high and consistent across cell types and species, is improved by balancing, but can be dramatically reduced at low read coverage.

**Figure 3.**
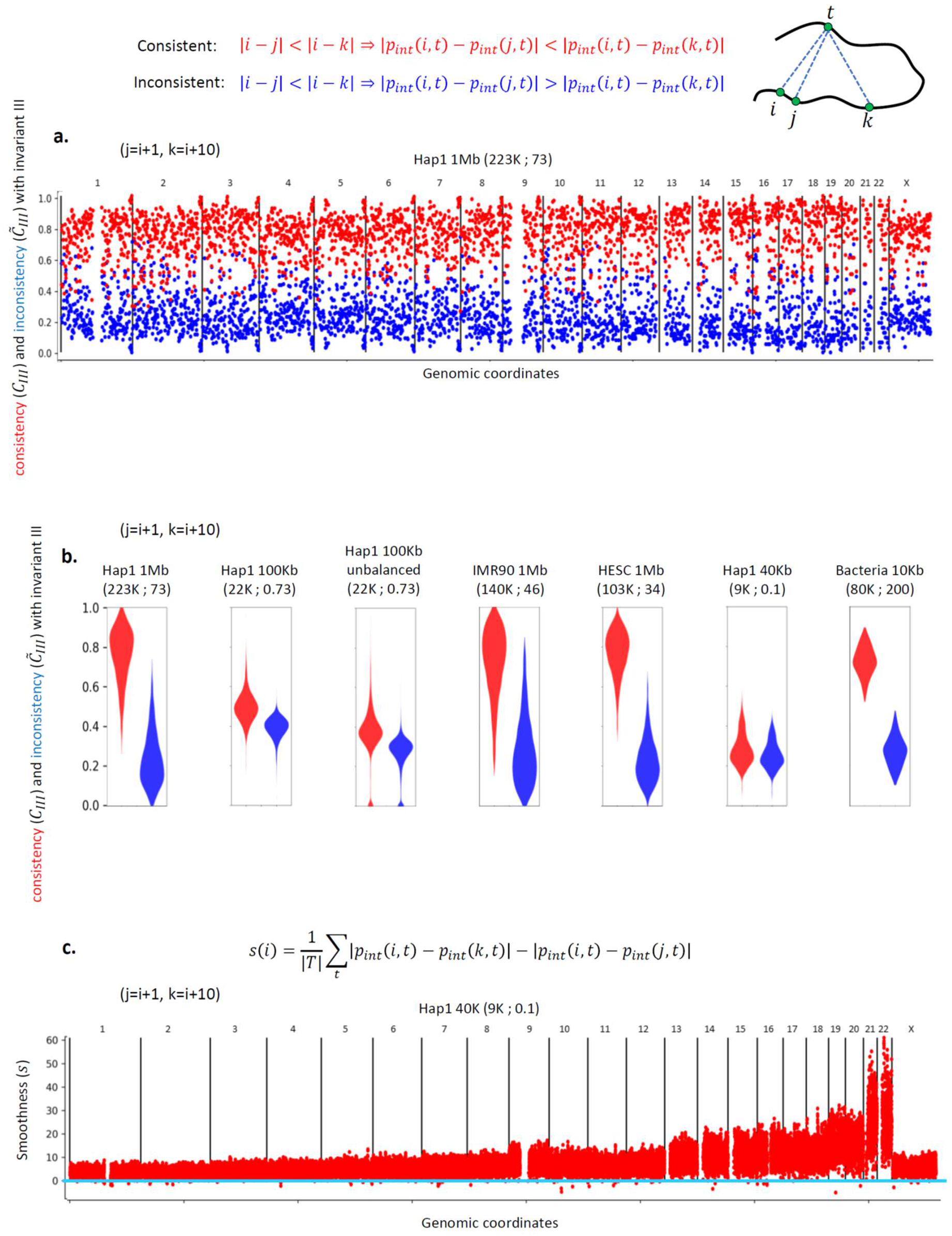
Genome-wide analysis of invariant pattern III (local interaction smoothness). (**a**) Genome-wide consistency (*C_III_* in red, see text for details) and inconsistency (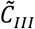 in blue) with invariant III for Hap1 Hi-C interaction map. (**b**) Violin plots showing the genome-wide distribution of *C_III_* and 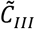 in various Hi-C interaction maps. (c) Genome-wide smoothness (s in red, see text for details) for Hap1 40K Hi-C interaction map. Cyan line indicates zero.

While the consistency with invariant III seemed to be low and potentially useless in low read coverage maps, we hypothesized that it may be useful to consider not only whether individual loci are consistent but also to what degree. To this end, we defined the *smoothness* of a locus as:

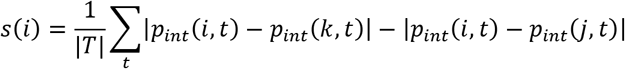

Where *P_int_*(*i,j*) is the interaction probability of loci *i,j*. Thus, a positive value of *s(i*) would indicate the intrachromosomal interactions of *j* are closer to those of *i* than the interactions of *k* are to those of *i*. We then calculated s genome-wide on a Hap1 40Kb resolution map, assuming positions *j = i + 1,k = i + 10* (units are bins) and that *t* are same-chromosome loci. Remarkably, we find that *s* values are overwhelmingly positive (0.999 of the bins are positive), in contrast to *C_III_* values. We suggest this difference is due to many inconsistent bins having low interaction probabilities. In light of these results, we suggest using *s* for practical applications such as detecting assembly errors and structural variation (see Evaluation section for an example of application in normal and extremely low-coverage interaction maps).

## 3. Hi-C-based scaffolding

Genome scaffolding is a sub-problem of genome assembly, which arises due to the inability to assemble the entire genome sequence during initial contig assembly. Genome scaffolding can be partitioned into 4 tasks: (1) Karyotyping: grouping contigs into chromosomes; (2) Contig ordering: determining the genomic order of contigs within each chromosome; (3) Contig positioning: determining the absolute position, or alternatively the gap size between contigs within each chromosome; (4) Contig orientation: determining the orientation of each contig with respect to other contigs within each chromosome. All Hi-C-based scaffolding methods use one or more of the invariant patterns. Invariant pattern I can indicate which contigs belong to the same chromosome, and can thus be used in karyotyping. Invariant pattern II can indicate which contigs within the same chromosome are closer than others, and can thus be used in contig ordering. By estimating the actual global distance decay interaction pattern, invariant pattern II can be used for contig positioning. Additionally, we can apply invariant pattern II to contigs that contain multiple restriction fragments and thus estimate contig orientation. Invariant pattern III indicates which contigs are near each other, but is less relevant to long-distance interactions. Invariant III can be useful for contig ordering and orientation, as well as error-detection (see Evaluation section).

Five Hi-C-based scaffolding methods have been developed previously: DNA Triangulation [23], LACHESIS [24], GRAAL [25], HiRise [40] and 3D DNA [29]. In general, current Hi-C-based scaffolding methods mainly use invariant patterns I and II, and use either probabilistic modelling approaches or graph-based approaches. Notably, the problem formulations suggested by all current methods do not admit optimal solution, and thus rely on heuristics. Of these, only DNA Triangulation, GRAAL and 3D DNA are actively maintained. All methods are provided via GitHub code repositories. Finally, we note that most methods do not try to resolve complex ploidy and repetitive regions.

Note that Hi-C-based scaffolding methods can either be applied directly to contigs or to mini-scaffolds produced by alternative technologies (e.g. long-insert libraries). To avoid confusion between input mini-scaffolds and the scaffolds constructed by Hi-C, we will refer to input contigs and mini-scaffolds simply as contigs since they are largely equivalent for the application of Hi-C-based assembly. In general, patterns in Hi-C become more pronounced with higher sequencing coverage and allow higher resolution, meaning large contigs will accumulate more reads and can be better positioned. In light of this, it is beneficial to aim for relatively large contigs in the input set (depends on the Hi-C coverage, but for current methods 100kb is a reasonable size to aim for). Finally, as we have observed that balancing improves consistency with all three invariant patterns, we recommend balancing Hi-C data prior to its usage in scaffolding.

In addition to the invariant Hi-C patterns, other interaction patterns appear in Hi-C maps. These patterns may be specific to the species, cell-type or cellular condition. These patterns include genomic compartments, TADs, point interactions, circular chromosomes and telomere/centromere clustering. While these patterns are typically not as strong as the invariant patterns, they can cause distortions and complicate Hi-C-based scaffolding. Indeed, in our analysis of invariant patterns we observe cell-type specific differences in some cases, as well as possible effects of circular chromosomes and telomere/centromere interactions. This potential link between errors in genome assembly and biological Hi-C patterns is important to keep in mind if the Hi-C is used to interpret 3D genome organization. For example, an assembly error in which a genomic region is left out may appear as if it is a TAD boundary, and vice versa, an actual TAD boundary may lead to prediction of a gap in the genome assembly. Thus, we recommend that biological features of 3D genome organization that are observed in Hi-C-scaffolded genomes should be interpreted cautiously (ideally, validated by orthogonal methods). An interesting notion is to try to experimentally eliminate the biological patterns leading to deviations from the invariant patterns. Indeed, this is the logic underlying Dovetail Genomics’ Chicago technology, which is based on extracting DNA and reconstituting it into chromatin before applying Hi-C[40]. However, it seems that this technique may have been replaced by standard Hi-C, possibly due to the loss of long-range interaction information due to DNA extraction.

## 4. DNA Triangulation

In the following section we overview our published Hi-C-based assembly method and provide practical guidelines for using it. DNA Triangulation [23] is maintained at https://github.com/KaplanLab/dna-triangulation.

### 4.1 General workflow

We recommend the following general workflow: (1) assemble initial set of contigs; (2) map Hi-C data to contigs; (3) partition contigs into bins (sub-contigs) of a set size, allowing for identification of problematic contigs, intrinsic evaluation of scaffolding, and contig orientation; (4) remove problematic bins and perform Hi-C correction using any available correction method (e.g. matrix balancing); (5) if chromosome number is unknown, estimate chromosome number using DNA Triangulation bootstrapped clustering method; (6) after chromosome number is chosen, partition contigs into chromosomes using clustering; (7) identify and remove problematic contigs based on clustering results; (8) scaffold each chromosome separately using probabilistic model; (9) remove problematic contigs; (10) evaluate results using orthogonal data and the invariant patterns. We next provide details on key points of this workflow.

### 4.2 Karyotyping

Karyotyping, including estimation of chromosome number, builds on invariant I and is performed using a bootstrapped variant of average-linkage hierarchical clustering. In order to estimate the number of clusters, we use an intrinsic property of the clustering process - clustering step length. We define a clustering step as the merging of two clusters in the clustering process. Each clustering step is associated with a merging distance, which is the distance between the two clusters which were merged. We then define the clustering step length as the difference between the merging distances of two consecutive steps. The clustering step length can be indicative of a stable partitioning [54]. Since hierarchical clustering in general, and the clustering step length in particular, may be sensitive to noise, we use a bootstrapped form of clustering by clustering a random fraction of the data and calculating the step length for each clustering run. Finally, we consider the average clustering step length over all runs, and find the maximal average cluster step. We estimate the number of chromosomes to be the number of clusters remaining at this point. In order to obtain the final partition into clusters, we finally cluster the entire dataset and partition by the number of clusters estimated from the bootstrapping.

With regards to estimation of chromosome number, we note that the default bootstrapping parameter of sampling 80% of the contigs can be adjusted when appropriate. For example, if it is suspected that there are small chromosomes containing few contigs, sampling 80% of the data may leave out significant portions of a chromosome and may thus cause underestimation of chromosome number. If this may be the case, we suggest increasing the portion of contigs sampled. In general, we suggest that it is better to overestimate the number of chromosomes in order to avoid chromosome misjoining. As long as there is some point in the clustering tree where the clusters are highly accurate, any clustering up to that stage would also be accurate, so even if we overestimate the number of chromosomes it should not affect cluster quality. Finally, we suggest using common sense when examining the average clustering step length profile - strong peaks may occasionally occur indicating a very low (2-3) or very high chromosome number (^~^1000). These can be excluded based on prior knowledge, and we suggest always considering secondary peaks in the relevant range. We recommend bootstrapping at least 100 times.

### 4.3 Chromosome scaffolding

Chromosome scaffolding is performed using a probabilistic model of a Hi-C interaction map assuming exponential distance decay. This method builds on invariant II, and in fact even makes a stronger assumption by specifying a parametric decay function. We then treat the positions of contigs and the decay slope as model parameters, and estimate these parameters using a maximum likelihood approach. Unfortunately, this optimization problem is non-convex (due to technical reasons we pose this as a minimization problem rather than a maximization problem) and cannot be solved optimally. Rather, we randomly initialize contig positions and gradually improve these using the L-BFGS algorithm until reaching a local optimum. This process is repeated several times (we recommend at least 10000) and the best (lowest) scoring solution is selected.

The predicted positions are given as numbers between zero and one, since we do not know in advance the size of the chromosome. However, by using the known distances between bins that belong to the same contig it is possible to infer the correct scale of the chromosome. This process is currently not automated in our code.

### 4.4 Contig partitioning

As detailed in the proposed workflow, contigs are partitioned into bins (sub-contigs) of a specified size (Figure 4). This is easy to perform, as most Hi-C matrices are binned by default. These bins are then scaffolded independently, without using the information regarding which contig they came from. This strategy has a few advantages:

1. <u>Contig and bin filtering:</u> Before karyotyping, after karyotyping, and after scaffolding, we can use the additional resolution provided by contig partitioning in order to evaluate the quality of the initial contigs. For example, if two bins belonging to the same contig are assigned to different chromosomes, this could potentially indicate an erroneous contig that could be filtered out or corrected. Additionally, misbehaving bins within a contig can be filtered out.
2. <u>Scaffolding evaluation:</u> If contigs are high quality, mis-assignment of bins that belong to the same contig may indicate a scaffolding error. In order to distinguish between the two scenarios, we recommend evaluating whether the invariants hold in each such inconsistency between contig assignment and scaffolding (see Evaluation section).
3. <u>Orientation inference:</u> In all Hi-C-based scaffolding methods, inference of contig orientation is based on the fact that contigs may contain several restriction fragments. Because the effective resolution of the experiment is higher than the contig size, it allows to use invariant II at sub-contig resolution, such that we expect the ends of contigs which are adjacent to interact strongly, and thus infer their orientation with respect to each other. This suggests an implicit approach to infer contig orientation using any Hi-C-based scaffolding method, by simply partitioning contigs into bins.

**Figure 4.**
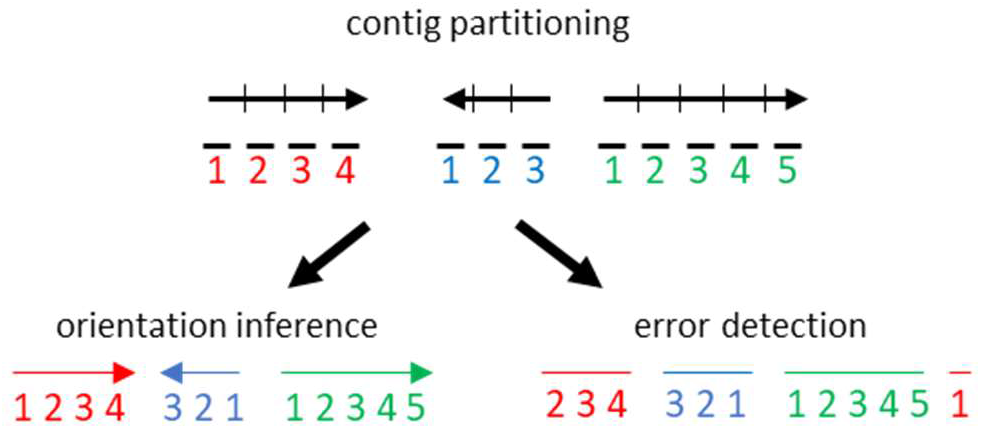
Contig partitioning.

The main disadvantage of contig partitioning is that predicting the positions of bins may be more difficult than the whole contigs, because each bin has fewer reads.

### 4.5 Parallelization

DNA Triangulation allows to massively parallelize the scaffolding process. For estimation of chromosome number, clustering is run several times on a random subset of data, providing robustness to outliers. This can be executed in a multithreaded mode (number of CPUs can be specified) in order to run these clusterings in parallel. The more computationally intensive task is the scaffolding of each chromosome, as it involves solving a non-convex optimization problem by gradually improving random starting positions. Thus, running more random initializations will likely improve the solution, so the major limitation is computational resources. Two parallelization options are provided for chromosome scaffolding: multithreading and HPCC, and these can act jointly. The number of random initializations and processors to use can be specified for each scaffolding run, and several such jobs can be run in parallel on a computation cluster. By keeping track of the optimization score of each job, we can finally select contig positions associated with the best (lowest) score.

## 5. Evaluation

An important aspect of any predictive algorithm in general and genome assembly methods in particular is being able to estimate the quality of the proposed solution without knowing what the correct solution is. In the case of Hi-C-based scaffolding, we propose a number of methods that can be used to evaluate solutions:

1. <u>Orthogonal data:</u> The first and most obvious way of validating a proposed Hi-C based scaffold is by using an independent technology, such as optical mapping or long-reads. Measurements made for validation purposes can be of relatively small scale, based on the principle that evaluating random parts of the proposed solution would give a reasonable estimate of the overall quality. However, these methods are costly and require the measurement of new data. Additionally, they mainly give an overview of solution quality rather than comprehensively detecting potential scaffolding errors.
2. <u>Contig partitioning:</u> As explained above, contig partitioning into bins can be used to evaluate scaffolding quality, by verifying that the bins belonging to the same contig are assigned to the same chromosome and that their predicted positions within the chromosome match their correct order.
3. <u>Invariants:</u> A comprehensive way of evaluating a scaffolding solution and comprehensively detecting potential errors is by finding deviations from the invariant patterns. It may seem futile to search for deviations from the invariants given that Hi-C-based scaffolding methods try to find a solution that is most consistent with these invariants. However, since these methods generally cannot find the optimal solutions, their proposed solutions can still deviate significantly from the invariants. Furthermore, invariant III (smoothness) is often not used explicitly by these methods, and can thus be used to detect potential scaffolding errors. In our experience, scaffolding errors are often easy to detect by using these principles, both visually and computationally. The ability to use intrinsic metrics to easily evaluate the quality of a scaffold (including one that was assembled by a different technology), is an important feature of Hi-C-based approaches.

To demonstrate the usefulness of invariant pattern III for error detection, we simulated a scaffolding error in human chromosome 4 in which two large genomic regions were misplaced (Figure 5a). We then calculated for both the original Hi-C map (Hap1 1Mb bins [48]) and the erroneous Hi-C the distance between every pair of adjacent bins. In line with our definition of smoothness, we define distance here as the mean absolute difference between the intrachromosomal interactions of bin *i* and those of bin *i + 1*. We then fitted a gaussian distribution to the adjacent bin distances, and used this distribution to calculate one-sided p-values for each bin to minimize noise and accentuate peaks. We observe that both the adjacent bin distance profile and the p-value profile of the mis-scaffolded interaction map clearly show three peaks corresponding to the misjoined loci. Next, since sequencing depth and resolution are important factors in Hi-C-based assembly, we asked whether the misjoined loci can be detected with significantly less reads. We simulated an extremely low-coverage scenario by sampling 100,000 reads genome-wide (^~^0.0001 of the original number of reads) from the Hap1 1Mb matrix. For the sampling, Hap1 1Mb read counts were normalized into probabilities and reads were sampled with replacement according to these probabilities. We then repeated the adjacent bin distance and p-value analysis. Albeit more noisy, we observe clear peaks that coincide with the error loci. Our results suggest that sparse Hi-C can be used to identify scaffolding errors, or similarly structural variation, using very small amounts of sequencing reads. In light of this, it might be useful in some scenarios to use Hi-C for scaffolding error detection even if the scaffolding itself does not use Hi-C.

**Figure 5.**
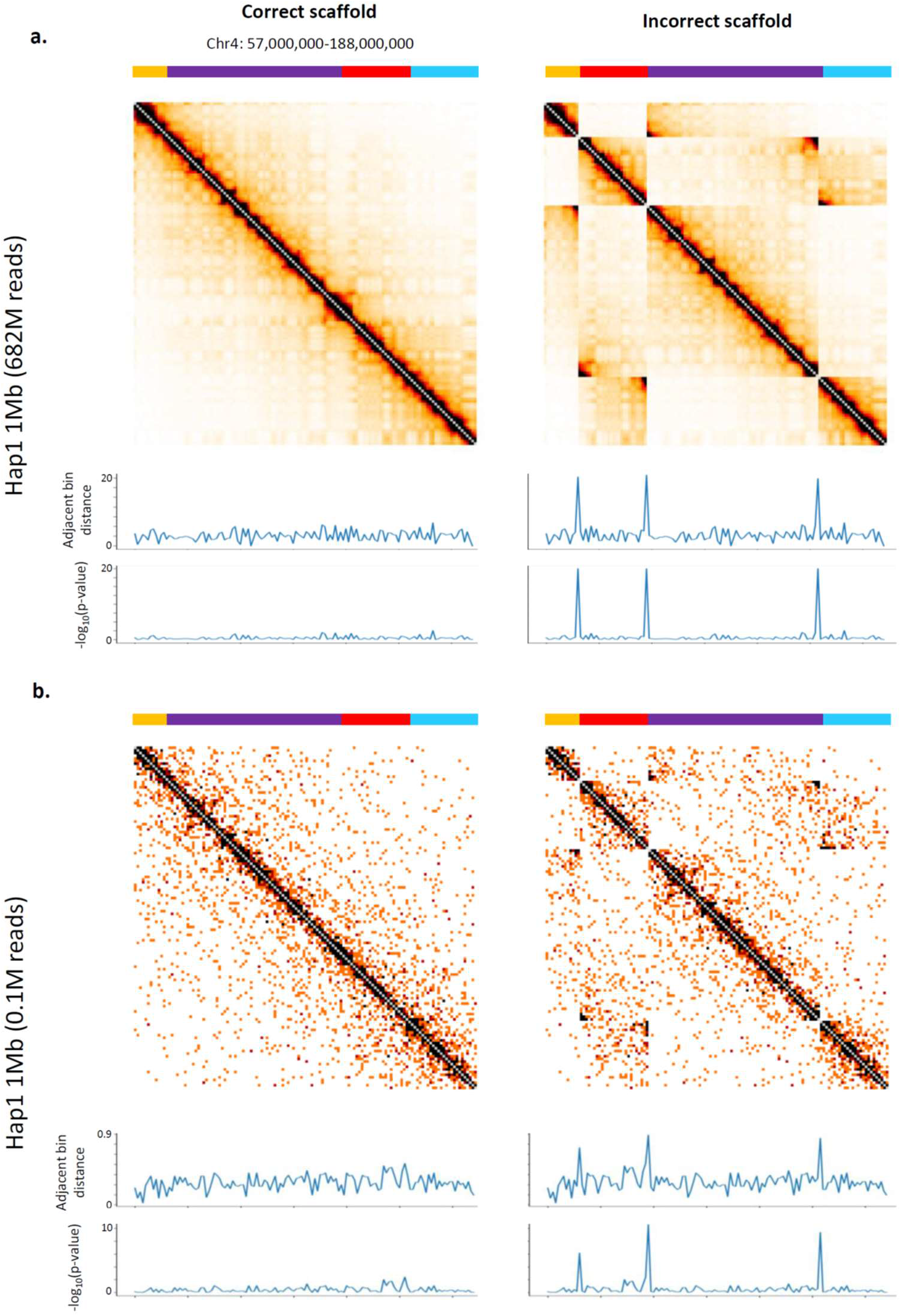
Detection of scaffolding errors or structural variation using invariant III. We simulated an incorrect scaffold by reorganizing a region of chromosome 4 (indicated by colors above Hi-C interaction maps). (**a**) Comparison of the correctly organized Hi-C map (Hap1 1Mb) with the incorrectly organized map. For each map we calculated the distance between every pair of adjacent bins. The incorrect map shows clear peaks at the error loci. We then fitted a gaussian distribution to the adjacent bin distances from the normal map, and used this distribution to calculate one-sided p-values for each bin as means of minimizing noise and accentuating peaks. P-values smaller than machine precision are. (**b**) In order to evaluate the performance of this metric in a highly sparse scenario, we used the Hap1 Hi-C map to sample 100,000 reads genome-wide (^~^0.0001 of the original number of reads) and repeated the analyses of (a). Albeit more noisy, we observe clear peaks that coincide with the error loci.

## 6. Discussion

High-quality genome sequences are central for understanding genome function on several levels. In spite of significant advances in genome assembly, there remains much room for improvement. *De novo* genome assembly, especially of large complex genomes, remains a major challenge. This is also the case for cancer genome assembly. While significant effort has gone into computational assembly methods based on short reads and long reads, only a handful of initial methods have been proposed for Hi-C-based assembly. Thus, one motivation for this work is to put current methods in context and stress that the full potential of Hi-C-based assembly approaches is not yet realized. In this paper we have quantitatively analyzed three invariant Hi-C patterns, in an attempt to better characterize their usefulness for Hi-C-based assembly. We suggest that simultaneously considering all three invariant patterns may lead to better Hi-C-based genome assembly methods. Furthermore, there are many opportunities for innovative computational approaches, theoretical research and introduction of novel concepts. A rigorous comparison of Hi-C-based approaches may also drive a newer generation of methods building on the respective strengths of individual techniques. Other challenges include incorporating Hi-C data in initial contig assembly, high-quality Hi-C-based assembly without intermediate technologies, hybrid Hi-C/long-read/long-insert scaffolding, and the identification of novel applications for Hi-C-based assembly.

## Acknowledgements

We thank all members of the Kaplan Lab. NK is supported by the Azrieli Faculty Fellows program and the Taub Fellows program.

